# A fast and objective hidden Markov modeling for accurate analysis of biophysical data with numerous states

**DOI:** 10.1101/2021.05.30.446337

**Authors:** Hanjin Liu, Tomohiro Shima

**Affiliations:** Department of Biological Sciences, Graduate School of Science, The University of Tokyo

## Abstract

The hidden Markov model (HMM) is widely used to analyze biophysical chronological data with discrete states, such as binding/detachment of biomolecules, protein/nucleotide conformational changes and step-like movement of single proteins. Despite its usefulness, classical HMM fitting has practical drawbacks that it requires the determination of the number of hidden states and fine initialization of many parameters before fitting. To overcome these drawbacks, several HMM pre-analyses have been reported, but do not provide enough accuracy when data have unknown kinetics and/or low signal-to-noise ratio. Therefore, in many cases, HMM fitting needs trial-and-error manual process that can impair the objectivity of the analysis. Moreover, for data composed of numerous hidden states, such as stepping data of cytoskeletal motors, there has been difficulty in HMM analysis because the large number of parameters were hardly properly initialized. Here, by combining a statistical step-finding method and the Gaussian mixture model clustering, we developed a new algorithm for more objective HMM analysis. Our algorithm can execute accurate state number estimation and parameter optimization with fully automated way. Simulation analysis demonstrated that our algorithm accurately fit both fast- and slow-transition trajectories. Compared with the previous method, the speed of our algorithm was 10–20 times faster for standard size data. Our algorithm also showed the accurate fit of the simulated motor-stepping data with more than 10 transition states, suggesting the applicability of the method to the data with numerous states. Furthermore, the algorithm is flexible enough to cope with cases where some kinetics are known in advance. Some available prior information, such as the dwell time of each state, can be integrated into the algorithm via two user-tunable parameters. In summary, our method enables fast, accurate and objective HMM analysis, and broadens the application range of HMM fitting that can provide more accurate interpretation of a wide variety of biophysical data.

## Introduction

Discrete chronological data are often seen in biophysical experiments, such as optical tweezer, single-molecule fluorescence resonance energy transfer (smFRET) and motor stepping. Since it is difficult to reveal the state transition from this type of data due to high-level noise, a large number of works have been exploring algorithms for appropriate data filtration (Chung *et al*., 1991; Watkins *et al*., 2005; Mckinney *et al*., 2006). Among them, Hidden Markov model (HMM) is a powerful tool for this purpose. HMM consists of a constant number of states, the probability distribution of observables, and a transition probability matrix, which indicates the probability of all possible transition between the state and state. By fitting a trajectory to HMM, one can interpret the trajectory as a series of stochastic transitions between states, and at the same time it can be easily translated to the expression of free energy changes. This translation allows scientists to elucidate the major reaction pathway of biomolecules, leading to gain a deeper understanding of the biophysical phenomenon. Furthermore, HMM fitting is performed by maximizing the probability product (or likelihood) to generate entire data using EM algorithm and state estimation using the Viterbi algorithm (Mukherjee *et al*., 2005), which are free of local optimal problem; thus, unlike smoothening-based filtering methods such as low-pass filter and median filter, the risk of ignoring rare state or to average out fast signal changes is lower.

The classical HMM requires the number of states to be determined beforehand. In some cases, the exact number is critical for the conclusion of the experiment. Nevertheless, it is determined simply by seeing histogram, which is affected by the bin width and accordingly causes a misinterpretation of the mechanism of biomolecules. Therefore, it is necessary to solve this problem by developing a method for state finding in a more objective manner. Previously, Mckinney *et al*. showed that using their program HaMMy, the number of states can be determined by repetitive

HMM fitting with a different number of states, followed by the determination of the most likely number of states by comparing BIC (Schwarz, 1978). However, the algorithm needs multiple (as many as 20) independent trajectories, which in some cases, is difficult or unfavorable due to the necessity of precise normalization procedures. Also, the algorithm turned out to be highly prone to overfit (Bronson *et al*., 2009). Therefore, the initial parameters, especially the number of states, should be determined in an objective, HMM-independent manner. To this end, it is natural to consider clustering methods such as X-means (Pelleg *et al*., 2000) and Gaussian mixture model (GMM), yet in the most cases the raw data trajectory should be filtered before clustering due to low signal-to-noise (S/N) ratio. Most methods available for filtering time series data, however, need at least one free parameter, and more importantly, it is difficult to assess the effect of parameter changes.

Kalafut-Visscher’s step-finding algorithm is first developed to achieve model-independent step finding of trajectories obtained from processive molecular motors (Kalafut *et al*., 2008). Since the number of steps and their positions is determined only by the mathematically established BIC minimization method, model independence is attractive for the above-mentioned purpose. We noticed that Kalafut-Visscher’s step-finding method can be applied not only to highly processive motor trajectories, but also to trajectories that contain many back steps as forward steps, like FRET efficiency obtained from smFRET experiments. This means that the step-finding algorithm is a promising tool for filtering HMM-type data before HMM fitting. However, its complete objectivity may conversely make it difficult to integrate information that the user knows a priori.

Here, we propose a step-finding-based HMM-fitting algorithm, sfHMM. We first modified Kalafut-Visscher’s method and derived an extended version of step-finding algorithm, which is based on the likelihood maximization instead of BIC minimization. By applying step finding and GMM clustering to HMM parameter initialization, we established a workflow that can conduct automatic HMM fitting with only two tunable parameters that are easy to interpret, and even these parameters can also be determined automatically. We demonstrated that, compared to HaMMy, sfHMM is far less computationally demanding and can find hidden states accurately. We also compared sfHMM with variational Bayesian method, _VB_FRET (Bronson *et al*., 2009) as a method for objective determination of the number of hidden states, and found that our method is not only 20–40 times faster for traces with standard time length, but also relatively tolerable to highly noisy traces and Poisson distributed traces. Moreover, sfHMM can also be applied to models with many states and sparse transition probability matrix, which were difficult to fit to HMM unless the initial parameters were precisely determined.

## Methods

We coded the algorithms for HMM fitting and simulation in Python scripts. For GMM clustering, we used scikit-learn library (version 0.23.1; Pedregosa *et al*., 2011). For HMM fitting and sample data generation, we used hmmlearn library (version 0.2.5). To compare the performance with that of _VB_FRET, the input/output format of the original _VB_FRET code is modified while the HMM-fitting part is kept untouched. To run _VB_FRET in Python, the main function of _VB_FRET was wrapped in Python scripts using matlab.engine, according to the official MATLAB website. The source code of sfHMM is available at https://github.com/hanjinliu/sfHMM.

### Extension of step-finding algorithm

To correctly fit raw data *d*_*t*_ to multi-step trajectory *θ*_*t*_, Kalafut *et al*. derived a BIC formula for detecting the correct step positions in *θ*_*t*_, assuming the standard deviation of the noise is constant (Kalafut *et al*., 2008). They showed that BIC minimization by successive optimal step addition is highly effective. When we applied the algorithm to trajectories generated from HMM, we found that in some cases, it ignores “bumping signals”, such as a forward step immediately followed by a backward step. To avoid underfitting, we tried to construct another criterion of judging step addition.

To this end, we introduced an average transition probability *p*_*sf*_ to likelihood function, which indicates the probability that *θ*_*t*_ will be a different value in the next frame of time. For the detailed derivation of formula, see Appendix A. In short, when *θ*_*t*_ has *k* steps, the resulting log-likelihood is written as following formula:

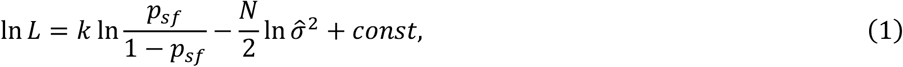

where 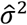 is maximum likelihood estimator for variance, and the last constant term is independent of step positions and the number. *k*-th step is added if ln *L* is larger than that with *k* − 1 steps. In our strategy overfitting will be prevented because in the case of *θ*_*t*_ with excessive steps, a small value *p*_*sf*_ is multiplied many times, which decreases the likelihood value. Thus, the first term in Eq. (1) can be interpreted as a penalty term like AIC and BIC.

In a specific case, maximizing log-likelihood ln *L* is mathematically equivalent to Kalafut and Visscher’s original BIC minimization because by substitution 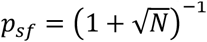, Eq. (1) becomes identical to the BIC formula:

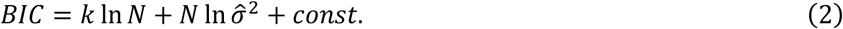

Thus, the likelihood max<u>im</u>ization model we derived is an extended concept of BIC minimization model. As can be seen from 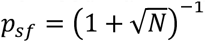, when the data size is large, BIC minimization model corresponds to likelihood maximization model with small *p*_*sf*_, leading to underfitting. This inconsistency occurred in BIC with different data size is understandable because BIC is an approximation form of Bayes factor assuming the data size is large. On the contrary, Eq. (1) is an accurate derivation, so that fitting will be consistent between datasets with different data size. Likelihood maximization model expanded the flexibility of step finding by adding one tunable parameter *p*_*sf*_ to Kalafut’s algorithm.

We also derived ln *L* assuming the signals follow Poisson distribution (Appendix B) because in some cases this assumption is more appropriate, such as single molecular experiments in which signals are proportional to photon counts. We demonstrated that step finding against traces following Poisson distribution also works (see Fig. 4B later in this article). We adopted this likelihood function for simulation data with Poisson distributed signals.

### Outline of sfHMM

The outline of sfHMM is shown in Fig. 1. First, the raw data *d*_*t*_ are fitted to multi-step trajectory *θ*_*t*_ that maximizes likelihood, by the step finding algorithm derived in 2.1. In sfHMM, one can set average transition probability *p*_*sf*_ manually, or directly apply Kalafut and Visscher’s method as a default setting. To set parameter *p*_*sf*_ is easy. If traces are sampled from fast transiting system like single molecule binding assay, i.e., the representative dwell time is short, one can set *p*_*sf*_ to, say, 0.1. In the case of slowly transiting system such as current profile obtained from nanopore, one can set *p*_*sf*_ to 0.01 or much smaller value.

**Fig. 1.**
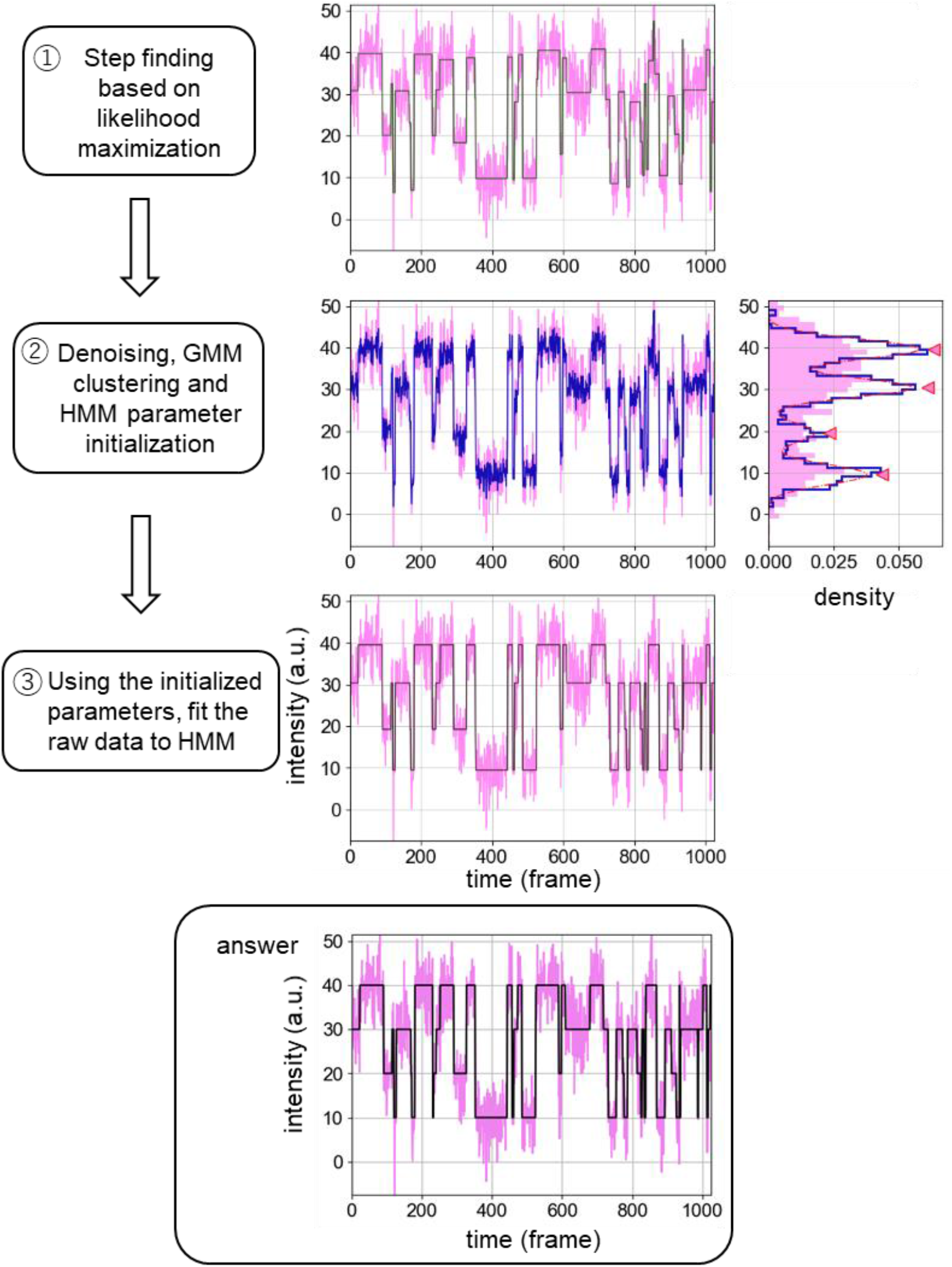
sfHMM fitting procedures. First, HMM sample data are fitted to multi-step trajectory *θ*_*t*_ by Kalafut-Visscher’s algorithm (1). Using *θ*_*t*_, raw data are denoised and subsequently applied to GMM clustering (2). These results are then used for HMM parameter initialization, and after that the raw data are fitted to HMM trajectory (3).

Second, the standard deviations between each set of adjacent changing points are decreased to *σ*_0_, which we call “denoising step”, by following formula:

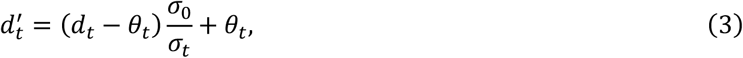

where *σ*_*t*_ indicates the standard deviation of raw data in the step which time *t* belongs to. If *σ*_0_ *is larger* than *σ*_*t*_ then the denoising step is skipped. *σ*_0_ is the second parameter that the user needs to predefine. However, it has the same unit as noise and signal intensity, which makes it much more intuitively understandable than parameters in other algorithms, such as window radius in median filter whose unit is time. Besides, as we will show later in 3.1, *σ*_0_ can be determined using *θ*_*t*_. Accordingly, unless the user knows the magnitude of signal changes between states, no assumption is needed beforehand.

The denoised data 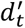 is then fitted to GMM function:

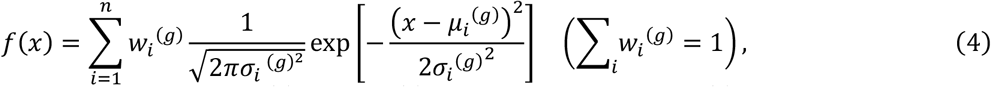

where the number of Gaussian *n*, the weight *w*_*i*_^*(g)*^, mean *μ*_*i*_^*(g)*^ and standard deviation *σ*_*i*_ ^*(g)*^ of the *i*-th Gaussian are determined by EM algorithm and BIC minimization. In this step, if any distance between two peaks is relatively small compared to *σ*_0_, it is plausible to say that this indicates overfitting. Hence, if any two adjacent peaks are closer than 1.5*σ*_0_, the result is considered to be insignificant. Any Gaussians with standard deviation less than 0.8*σ*_0_ was also ignored, since theoretically the resulting standard deviation of the denoised data should be ≳ *σ*_*0*_.

*All t*he parameters determined above are then utilized to initialize the parameters used for HMM fitting: initial probability distribution Π_*i*_, transition probability matrix *W*_*ij*_, mean values *μ*_*i*_ and variances *v*_*i*_, where suffix *i* denotes the *i*-th state. Since it is natural to assume that the peak positions of the histogram of denoised data are close to the real mean values of the hidden state, HMM’s mean values are simply initialized by GMM’s mean values:

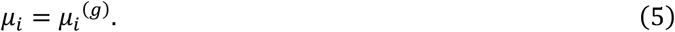

Before initializing *W*_*ij*_ and *v*_*i*_, all the raw data points are assigned to any of the *n* states. This can be attributed to the problem of inferring hidden state trajectory *s*_*t*_ without using conventional HMM fitting methods. If *θ*_*t*_ is well fitted, then *θ*_*t*_ should look similar to *s*_*t*_. Therefore, *s*_*t*_ is set to the state that *θ*_*t*_ is the most likely sampled according to GMM:

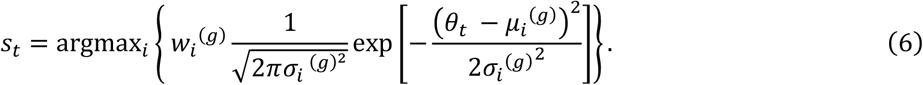

With the hidden state trajectory *s*_*t*_, transition probability matrix is initialized by empirical distribution:

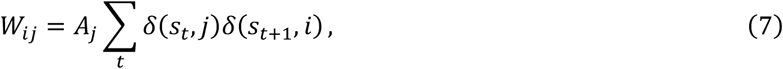

where *A*_*j*_ is normalization constants to satisfy ∑_*j*_ *W*_*ij*_ *= 1*, and *δ(x, y)* is delta function defined as *δ(x, x)* = 1 and *δ(x, y*) = 0 when *x ≠ y*. Variance of *i*-th state is initialized by calculating the sample variance of all the data points belong to *i*-th state.

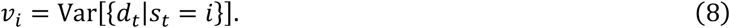

Finally, initial probability distribution is initialized using the rest of the parameters:

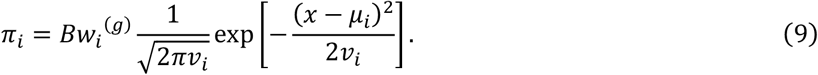

where *B* is also a normalization constant.

## Result

### 1. *σ*_0_ determination method and its evaluation

Although decreasing *σ*_0_ leads to more expectation to find states that are buried in random noise, it also causes over-estimation of the number of states. To correctly separate peaks using GMM clustering, the random noise should at least be cut off to 20% of the distance between two peaks (Fig. S1). Therefore, if the signal changes upon transition are known, one can minimize the risk of over-estimation while keeping the risk of under-estimation low by setting *σ*_0_ to 20% of the signal change. We noticed that for various types of trajectories, the signal change can well be estimated from the step size list of *θ*_*t*_, simply using the 25th percentile of the list (Fig. S2). Therefore, after the step size list of *θ*_*t*_is given, we set *σ*_0_ to 20% of the 25th percentile of step size list.

To evaluate the validity of this *σ*_0_ determination method, HMM trajectories were generated with various numbers of data and Gaussian noises, and the under/over-estimation frequencies were counted (Fig. 2). Because the whole algorithm is independent of the unit of observables for Gaussian distributed traces, we normalized the signal change between each set of adjacent states to 1. To simplify the model, all the hidden states were considered equivalent, and the transition probability matrix is assumed to satisfy *W*_*ii*_ = 1 − q and *W*_*ij*_ = q/(*n* − 1), where *n* denotes the number of states.

**Fig. 2.**
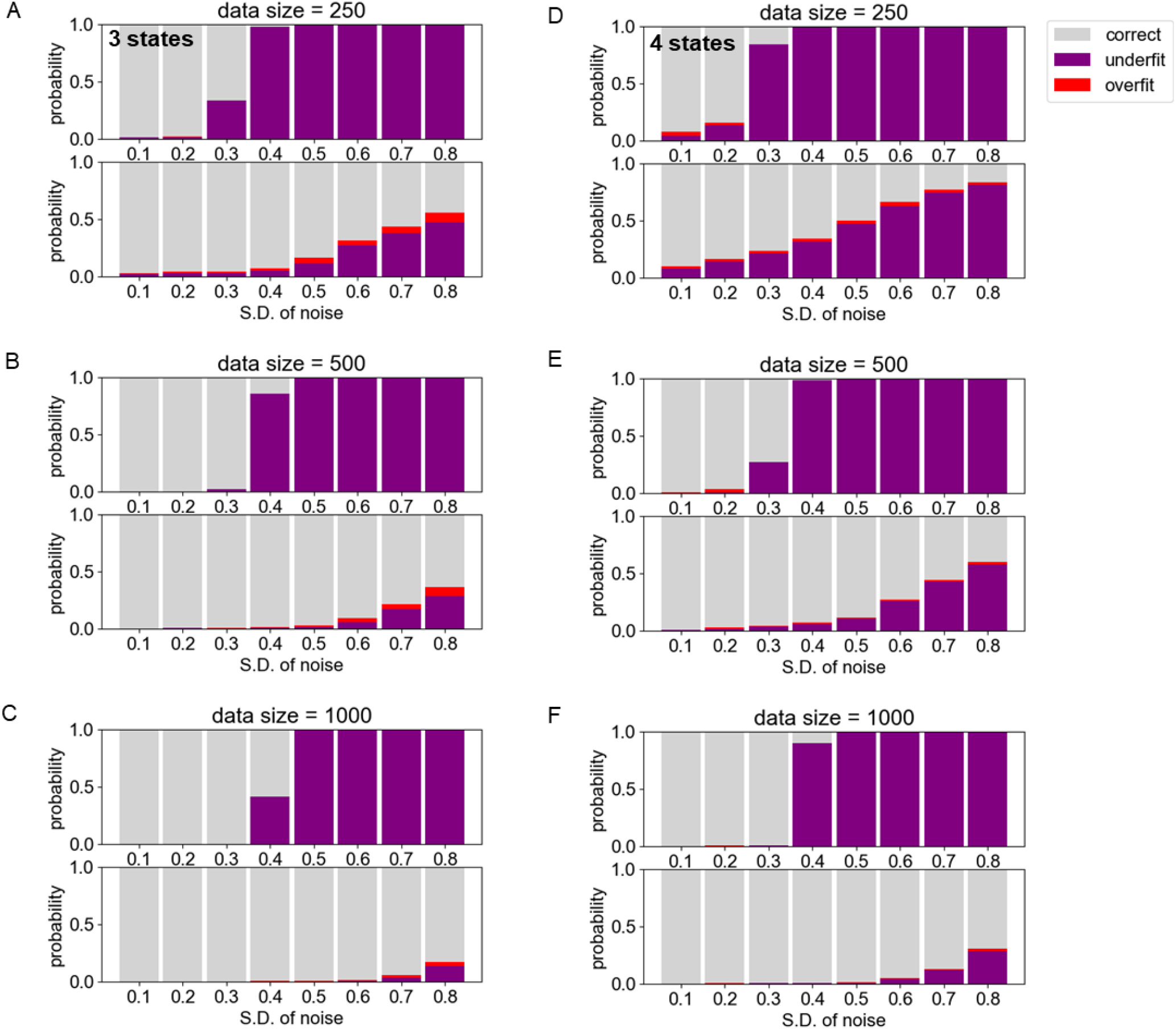
The probabilities of over-estimation (red) and under-estimation (purple) of the number of hidden states by GMM clustering, with varieties of total number of frames and noises. Three-state model (A-C) and four-state model (D-F) are simulated. For each figure, the upper bar graphs are the results obtained by directly clustering the raw data, and the lower ones are the clustering results with data denoised by our method using default *p*_*sf*_. The real mean values were set to {1, 2, 3} for three-state model, or to {1, 2, 3, 4} for four-state model. For simplicity, all the diagonal components of transition probability matrix were set to 0.95 and all the non-diagonal components were assumed to be same. The standard deviation (S.D.) of noise is independent of the hidden state and is set to the same value.

As expected from the result of Fig. S1, real number of states can hardly be found from raw data trajectory with noise more than 0.3. On the contrary, after the denoising step, the number of states were almost correctly estimated when the noise is ≤ 0.5, and in some conditions, 50% correct even the noise reached 0.8. This outcome is even better than that of low-pass filtering, which is a simple method to eliminate noise, or Chung-Kennedy filtering, which is widely used for filtering data with discrete states (Fig. S3), even though the user-defining parameters for these filtering methods are carefully chosen. It is noteworthy that low-pass and Chung-Kennedy filterings are highly prone to overfit when noise is weak. This means that excessive filtering will lead to detection of “pseudo peaks”.

We concluded that in the standard three- or four-state transition, state number can be well estimated by our method for the most part of the trajectories with noise as much as 0.5-0.6.

### 2. Performance of sfHMM

To compare the result of sfHMM with that of other methods, 1000 trajectories were generated and filtered with a diverse of method, and the mean absolute error (MAE) values were calculated for each result (Fig. 3A). Examples of filtering results are shown in Fig. 3B-E. As can be seen in Fig. 3A, sfHMM worked better than widely used low-pass filtering and Chung-Kennedy filtering. This is because low-pass filter and Chung-Kennedy filter uses a constant cutoff frequency or window radius respectively to smoothen data, so that they have a risk of smoothing out steps narrower than the lengths or detecting step-like parts within a single step (Fig. 3D, E). We also analyzed the sample trajectories only using HMM fitting. Here, the correct number of hidden states *n* = 4 were given, and the transition probability matrix was roughly initialized to *W*_*ii*_ = 0.1 and *W*_*ij*_ = 0.1/3, which corresponds to average dwell time being ∼10 frames. Mean values were initialized using k-means clustering method, and variance of each state were set equal to the variance of the entire data. As shown in Fig. 3, MAE of our algorithm almost reached the lower limit of HMM fitting, suggesting that all the uncertainties except for unavoidable error during HMM fitting were solved by our HMM parameter initializing method.

**Fig. 3.**
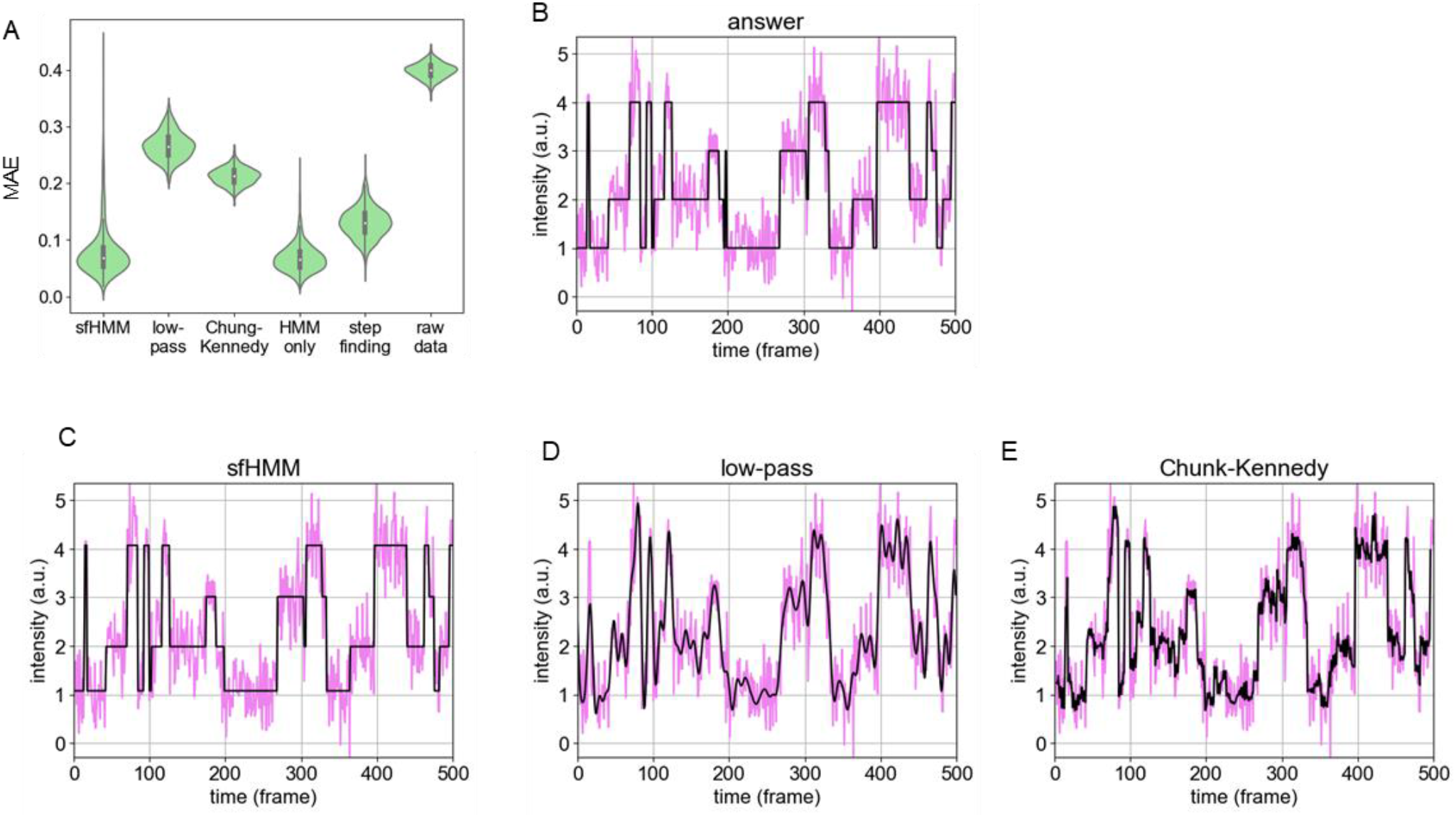
Comparison between filtering methods. Four-state, 500-frame HMM data were sampled 1000 times. (A) MAE distribution of each algorithm is shown using violin plot. The inner bar plot denotes median, 25% percentile and 75% percentile of each set of simulation results. MAE is defined as 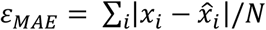, given that the real values *x*_*i*_ are estimated to be 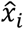. “HMM only” denotes that raw data were directly fitted to HMM using four-state model, the correct number of hidden states. “Step finding” denotes the intermediate result in sfHMM. (B-E) An example of sample data and its fitting or filtering results are plotted in the same way: (B) the real trajectory; (C) inferred real trajectory using sfHMM; (D) low-pass filtering with cutoff frequency = 0.1 per frame; (E) Chung-Kennedy filter with 3-frame window radius and weight parameter 5.

Subsequently, to see robustness of algorithm, we applied sfHMM to trajectories generated from different models (Fig. 4). Again, the results of the simple HMM fitting using correct number of states were also shown for comparison. First, we used several transition probability matrices *W* to express different dwell times *τ*. We suppose one stable state and two unstable states (*τ* = 5, 10, 15, 20) in a system, and defined *W* as:

**Fig. 4.**
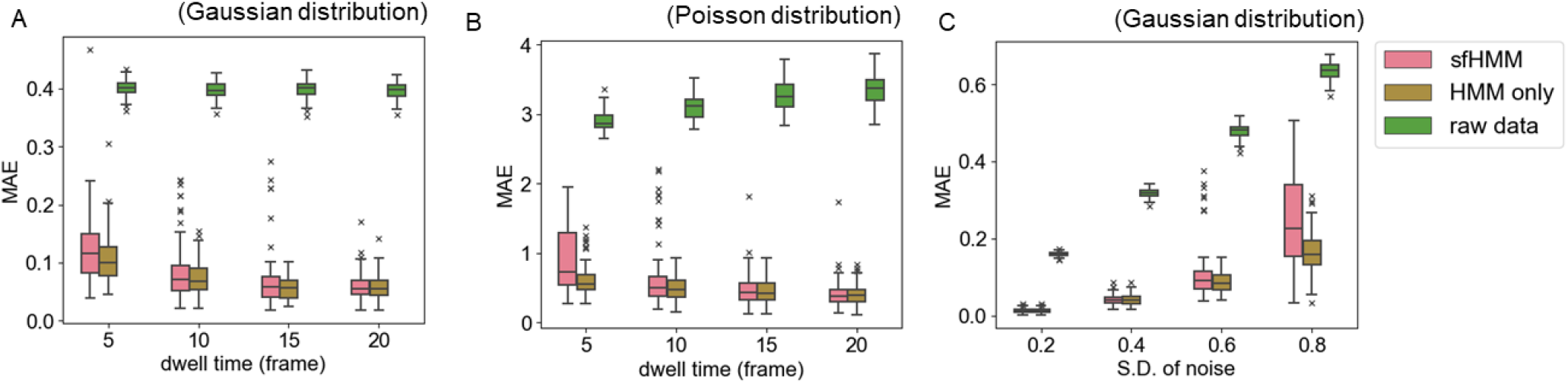
Three-state, 500-frame HMM data were sampled 100 times assuming Gauss distributed signals with mean values {1, 2, 3} (A, C) or Poisson distributed signals with mean values {10, 20, 30} (B). Sample datasets were fitted by sfHMM, or directly by HMM fitting algorithm using the correct number of states. Boxplot of MAE is shown against parameter changed during simulation. The MAEs of raw data are also shown for comparison. (A) The average dwell time *τ* of unstable states is changed from 5 frames to 20 frames, which corresponds to diagonal components of transition probability matrix ranging from ∼0.82 to ∼0.95. Standard deviation of noise is fix in 0.5. Average transition probability *p*_*sf*_ is set to 0.08 for *τ* = 5, 10 and to 0.04 otherwise. (B) Same condition as A, but the noise follows Poisson distribution. *p*_*sf*_ is set to the same value as (A). (C) Standard deviation of Gaussian noise is changed from 0.2 to 0.8, where the signal change between adjacent states is set to 1. All the diagonal components of transition probability matrix are set to 0.95, which corresponds to average dwell time ∼20 frames. The default *p*_*sf*_ is used in sfHMM.

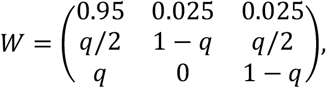

where *q* is determine by *q* = 1 − exp(−1/τ), namely, *q* ≈ 0.18, 0.10, 0.064, 0.049. We roughly set average transition probability *p*_*sf*_ assuming the real dwell time is unknown: *p*_*sf*_ = 0.08 for fast transition (*τ* = 5, 10) and *p*_*sf*_ = 0.04 for slow transition (*τ* = 15, 20).

Because step finding has a risk to ignore short time lived bump, our algorithm failed to accurately fit more than five simulated trajectories out of 100 when the average dwell time of unstable states is 10 frames or shorter (Fig. 4A). If it is longer than that, the average fitting errors were almost the same as the lower limit of HMM fitting. sfHMM performed similarly well for Poisson distributed signals (Fig. 4B). Strictly, average transition probability equals to the summation of non-diagonal components of transition probability matrix. To correctly detect unstable states, we should have set to *p*_*sf*_ = q. Therefore, our method can accurately carry out HMM fitting even if *p*_*sf*_ cannot be accurately determined in advance.

With larger Gaussian noise, the average MAE of sfHMM results also increased (Fig. 4C). This does not necessarily mean sfHMM failed, because the average MAE also increased when only HMM fitting is used. Taking Fig. 2B into consideration, sfHMM can estimate optimal HMM parameters unless the number of states were wrongly determined, which corresponds to the occasional MAE increase in Fig. 4C.

In summary, our method can be used in wide range of trajectories with different distribution and S/N ratio.

### 3. A method of combining multiple trajectories from different molecules

We have demonstrated that sfHMM is incredibly useful for wide range of data. However, the data sizes we can get are usually limited in standard single molecule experiments because of photobleaching or dissociation from the glass surface. Therefore, practically we sometimes need to combine multiple trajectories obtained from distinct molecules.

We hypothesized that our algorithm can easily be applied for this purpose by following three procedures:

1. Link all the step finding results to determine overall *σ*_0_.
2. Denoise every dataset using overall *σ*_0_ and accumulate all the denoised data to fit to GMM function.
3. Initialize HMM parameter similarly as referred in 2. but using all the hidden state trajectories *s*_*t*_. Log-probability is calculated along all the trajectories during EM algorithm so that the “consensus” HMM parameters are determined.

To test effectiveness of this method, a set of sample data with small data size were generated and fitted. As shown in Fig. 5A-B, this strategy worked well with small-size data. To compare the performance of this algorithm with that of Chung-Kennedy filtering, we repeated simulation 100 times, and found that MAE values of our method were better (Fig. 5C) except for the occasional MAE increase caused by underfit. This result proved the advantage of our algorithm over classical filtering method. We also tested this strategy to see whether it works in the case of photobleaching assay or experiments using motor proteins. Again, sfHMM successfully found the steps in both cases (Fig. S4). Note that in the photobleaching assay, it takes little time to pass through the first few steps, which makes step finding difficult. Likewise, it is difficult to correctly fit to the short-stayed position of motor trajectory. Nevertheless, by gathering three trajectories, sfHMM successfully detected almost all the steps.

**Fig. 5.**
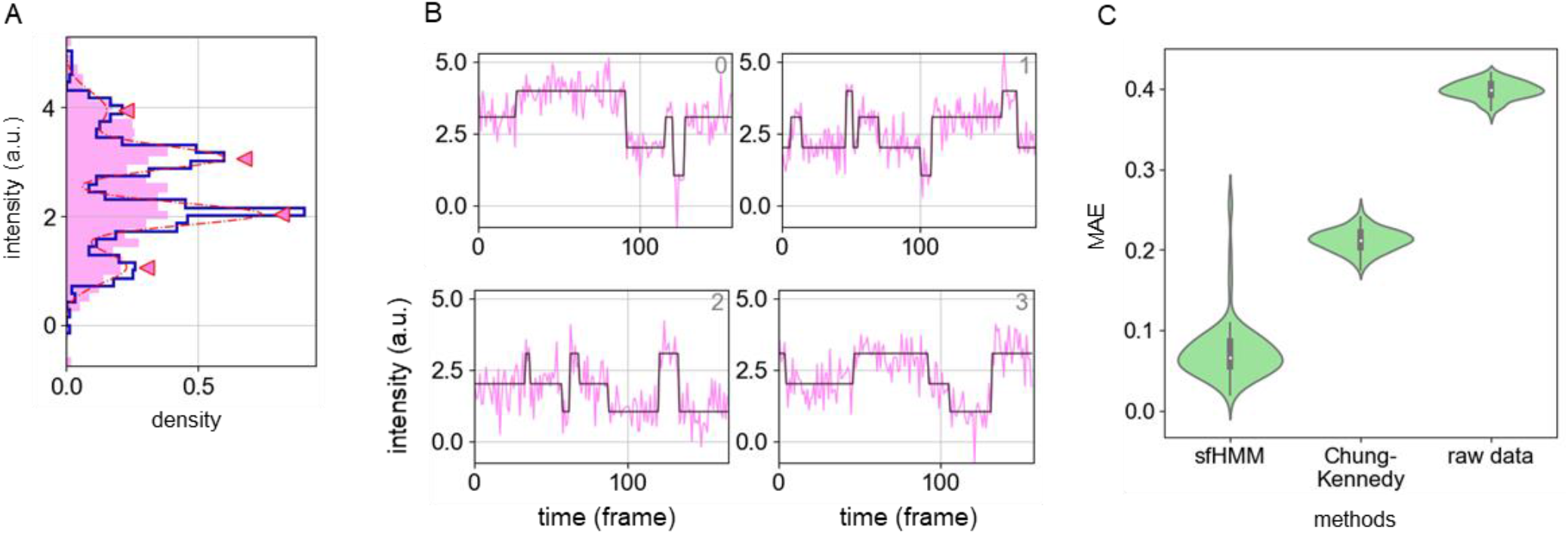
Four-state sample data are repetitively generated and subsequently four trajectories are combined before HMM fitting. Since the observed time length cannot be guaranteed identical among molecules due to photobleaching or other experimental artifacts, each trajectory have different data sizes, distributed uniformly between 70 and 230. (A, B) An example of the sfHMM result, by combining four datasets. (C) Simulations were carried out 100 times and MAE is plotted using violin plot. The results of Chung-Kennedy filtering with 3-frame window radius and weight parameter 5 are shown for comparison.

### 4. Comparison with an existing method _VB_FRET

_VB_FRET (Bronson *et al*., 2009) is also a method that can infer HMM without user-dependent input. For example, if one wants to know the best model among one-state model to a five-state model, raw trajectory is repetitively fit to HMM and a conditional probability called evidence is then calculated for each model. The model that maximizes evidence is finally selected.

To compare the performance of our algorithm to that of _VB_FRET, again sample data were repetitively generated, and each trajectory is processed using both algorithms. First, as in Fig. 4, we evaluated the fitting result of each algorithm by MAE, using trajectories with different dwell times and noises. Intuitively, _VB_FRET is supposed to return better fitting results for fast transiting trajectories because the hidden states are inferred only by HMM fitting, unlike one-by-one step-finding algorithm that is at risk of ignoring bumping signals. However, the difference between the two methods turned out to be trivial for Gaussian distributed signals (Fig. 6A) and sfHMM fitted even better for Poisson distributed signals (Fig. 6B). sfHMM may be able to deal with fast bumping signals because bumping happens many times throughout trajectories. During step finding, when a signal change point is detected at one side of the bump, the *χ*^*2*^ value of unstable states decreases, while that of stable state increases. Thereby, if *χ*^*2*^ value of unstable states will decrease enough, or in other words, if the hidden state was in one of two unstable states for long time enough, the total *χ*^*2*^ value will decrease and the likelihood will become larger.

**Fig. 6.**
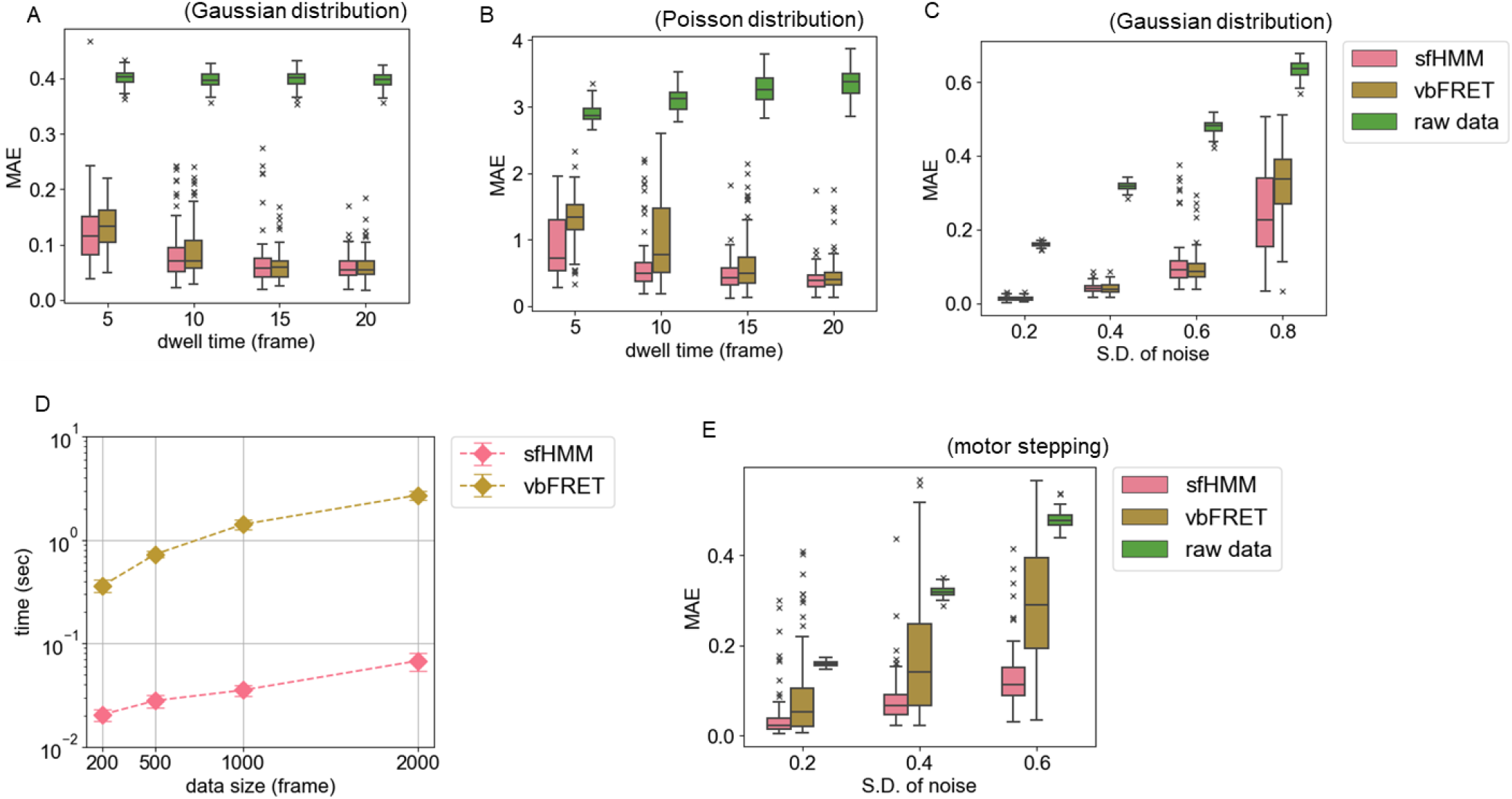
Comparison between sfHMM and _VB_FRET. Three-state, 500-frame HMM data were sampled 100 times. (A-C) Simulation conditions are same as Fig. 4A-C, except for _VB_FRET is used in place of “HMM only”. (D) Processing times (mean±S.D.) that sfHMM and _VB_FRET took to fit data with different size. To compare in the same condition, model selection was carried out from two-state model to five-state model for both algorithms. To confirm fitting precision, parameter initialization was repeated five times in _VB_FRET for each trajectory. To exclude irrelevant processing time, time measurement is executed within the MATLAB code of _VB_FRET. (E) HMM fitting of motor stepping trajectories. Default *p*_*sf*_ is used in sfHMM. Molecular motor is assumed to have forward step probability 0.02 and backward step probability 0.01.

Two algorithms were able to fit similarly well with noise under 0.6, but if the noise got as high as 0.8, our method showed relatively lower MAE (Fig. 6C). This may partly own to a high level of noise-tolerance of step-finding algorithm. Also, the difficulty of fitting noisy data using _VB_FRET may be an adverse effect of comprehensive search for the best hidden state. Before HMM parameters almost converged to the real values, it is hard to separate noise from fast state transition, especially when the signal change of noise approached that of state transition.

Besides, we noticed that _VB_FRET is considerably time-consuming (Fig. 6D), taking 20–40 times longer compared to sfHMM. Although sfHMM and _VB_FRET are written in different languages (Python and MATLAB respectively), this conclusion would not change because _VB_FRET must repeat HMM fitting, which is computationally much more expensive than GMM clustering, the rate-limiting step in sfHMM. Indeed, by measuring the processing time of HMM fitting in Python, we estimated that _VB_FRET would take ∼2.4 seconds with 500-frame data, which are almost the same because of MATLAB-coded _VB_FRET (1–2 seconds). Although it is possible to save time by decreasing the repetition in _VB_FRET, the accuracy of _VB_FRET became exceedingly low in that way.

We also tested whether two algorithms can be applied to motor-stepping trajectories. In terms of HMM, motor-stepping trajectories have many hidden states and some of them exist for only a short time, so that HMM fitting is highly prone to underfit the trajectories. By comparing MAE, obviously _VB_FRET results in higher error (Fig. 6E). To examine in more detail, we evaluated fitting results of two algorithms by the number of states found, which corresponds to the number of positions molecular motor bound to, and by the number of transitions, which corresponds to the number of stepping movements molecular motors did (Fig. S5A). When the standard deviation of the noise is 0.2, sfHMM was able to find the correct number of states from 48 trajectories out of 100, while this number decreases to 26 for _VB_FRET (Fig. S5B). Other conditions displayed the same trend. When we focused on the number of found steps, the difference was more obvious (Fig. S5C). When the standard deviation of the noise is 0.2, sfHMM was able to detect the number of steps with an error of less than two steps from 64 trajectories out of 100, while the number decreases to 41 for _VB_FRET, partly because of under-estimation of the number of states.

In summary, for analyses of Poisson distributed HMM trajectories, motor-stepping trajectories and a large number of trajectories, sfHMM should be the better choice.

## Discussion

Our program sfHMM enabled HMM fitting to apply to various trajectories with or without intuitively tunable parameter inputs. We demonstrated that, to correctly estimate the number of hidden states, denoising based on step-finding algorithm is even better than that usually used low-pass filter or Chung-Kennedy filter with carefully determined input parameters (Fig. 2, Fig. S3). Compared to other filtering methods, sfHMM predicted the hidden state much more accurately, and almost reached the limit of HMM-fitting accuracy (Fig. 3). It also worked with various dwell times, S/N ratios, and distributions (Fig. 4). Recently, smFRET is widely used to analyze the intermediate state of protein conformation (Osuka *et al*., 2018; Stella *et al*., 2018). If trajectories are analyzed using the filtering method listed above, one must pay attention to pseudo-peaks because overfitting frequently occurs when user dependent parameters are not appropriately chosen. In this sense, to accurately quantify the contribution of the newly emerged states, sfHMM-based approach will be useful.

In the case of the demand for analyzing data acquired from a short time, we successfully combined multiple trajectories to fit to such data. This strategy worked with only four trajectories, ∼600 data points (Fig. 5). It was also useful when applied to HMM system with a sparse transition probability matrix, such as photobleaching and motor stepping (Fig. S4). Both situations are not suited to direct HMM fitting because of the parameter initialization problem. It can be said that sfHMM broadened the application range of HMM fitting.

By comparing with _VB_FRET, we found that although two algorithms performed equally well in most cases, sfHMM was relatively noise-tolerant (Fig. 6C), applicable to motor-stepping trajectories (Fig. 6E, Fig. S5) and requires much less calculation time (Fig. 6D). We also noticed that if all states are equally unstable, that is, all the diagonal components of transition probability matrix are small, _VB_FRET performed much better than sfHMM (Fig. S6). This situation will be comparatively rare because even if free energy values differ only for *k*_*B*_*T* two distinct states, one of the average dwell times will be around three times longer than the other. As long as signal acquisition rate of the observation system is fast enough to catch the short-dwelled steps, obtained trajectories should have at least one stable state, as assumed in Fig. 4. However, unless one is sure about the underlying kinetics, it is important to test HMM fitting in different ways and discuss comprehensively.

Because sfHMM consists of separate procedures: step-finding-based denoising, GMM clustering, HMM parameter initialization and HMM fitting, sfHMM has high potential to customize the algorithm flexibly. For example, it is possible to widen the usage of sfHMM by replacing HMM analysis with causal state model analysis (Kelly *et al*., 2012). Although causal state model can distinguish different hidden states even if signal intensities degenerate, the original algorithm imposed stringent requirements such as high S/N ratio and sufficient data size. Both requirements can be solved by implementing the causal states modeling algorithm into sfHMM workflow: cutting off noise in the denoising step and increasing data size by combining trajectories in the way shown in 3.3. Another example is to make sfHMM specialized in photobleaching analysis. Because trajectories obtained from photobleaching experiments contain different variances, it is inappropriate to apply step-finding algorithm assuming a Gaussian distribution with constant variance. Although we also derived an algorithm of the Poisson distribution model, fluorescence intensities must be translated to photon counts and basal noise may prevent accurate step finding. Previous works (Chen *et al*., 2014; Tsekouras *et al*., 2016) have provided step-finding algorithms optimized for photobleaching assay, thus these methods can be an alternative for the step-finding algorithm we used.

We have simulated Poisson distributed trajectories by setting the photon count change upon state transitions to 10 (Fig. 4B, Fig. 6B). In some cases, however, less photon may be observed due to the fast image acquisition rate. Unlike the Gaussian distribution, Poisson distribution is unscalable, so that fitting universality before and after scale conversion is not guaranteed mathematically. Indeed, when the photon count change is set to 3, sfHMM failed to detect hidden states (Fig. S7). Appa<u>r</u>ently, it<u>is</u> due to the difference of S/N ratio. When mean values are *{λ, 2λ, 3λ}* then the s<u>tan</u>dard <u>dev</u>iations are 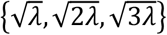, thus S/N ratio between the second and the third states is calculated to be 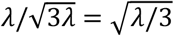. It is difficult to infer the hidden state trajectory accurately when *λ* is small. We also tried the previous Bayesian step detection method (Ensign *et al*., 2010), but still, it could not deal with the noisy trajectory. It seems necessary to ensure large *λ* by high laser power or long exposure time.

If the noise is not white, one should use step-finding algorithm other than Kalafut-Visscher’s approach, since it is extremely sensitive to correlated noise (Little *et al*., 2011). Correlated noise also makes GMM clustering harder because the signal will be highly biased depending on the previous state, which distorts the resultant histogram. The dynamics of biomolecules are correlated with principle since the positions of atoms are contiguous to time. To successfully apply sfHMM, one must set the time bin long enough compared to the time scale of dynamics, which causes signal changes, as well as short enough compared to the average dwell times.

Kalafut-Visscher’s method is first introduced for fitting to motor-stepping trajectories. However, as described in their paper, it is prone to overfit at low noise-level trajectories. Our simulated data also showed overfitted steps (Fig. 1). It seems that this type of step-finding method tends to recognize short-lived “halfway steps” within true steps. The reason is that a small number of data may accidentally have much lower standard deviation than expected, so that the *χ*^*2*^ value in the likelihood function largely decreases. Although this error is inevitable, it can be eliminated after GMM clustering, as can be seen by comparing the multi-step fitting result and the HMM-fitting result in Fig. 1. Therefore, low-overfitting-rate step finding of motor trajectories can also be achieved by applying our algorithm. This is important for some cases, such as revealing dynamics of a molecular motor dynein. Dynein occasionally skips a step, resulting in a mixture of 8-nm and 16-nm steps (Kinoshita *et al*., 2018), so that it is difficult to reasonably determine whether each step is 8-nm or 16-nm without HMM fitting. However, sfHMM still has a high probability to fail to detect all the states in a single motor-stepping trajectory (Fig. 6E, Fig. S5). This problem may be solved by considering constraints such as the periodicity of step positions in the future.

## Acknowledgement

This work was supported by Japan Society for the Promotion of Science KAKENHI (19H05379 and 21H00387) to T.S. We thank the members of the Uemura laboratory for valuable discussions.

## Author Contributions

Conceptualization, H.L. and T.S.; Analysis, H.L.; Writing, H.L. and T.S.

---

**Fig. S1.**
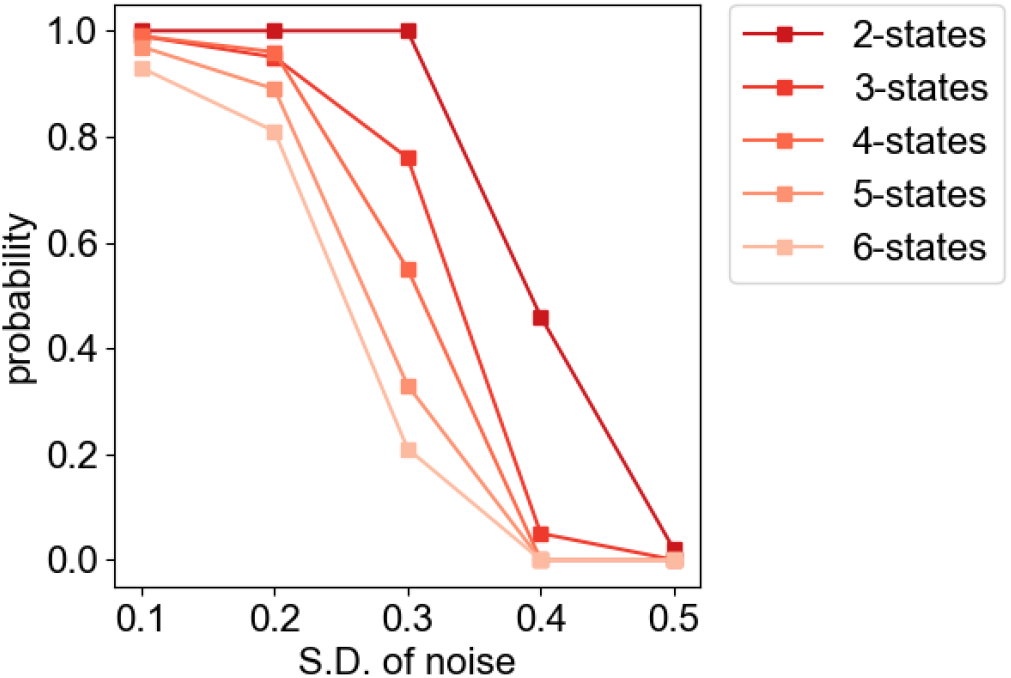
Standard deviation of Gaussian noise that GMM fitting tolerates. Simulations were run 100 times for each condition, and the probabilities that the number of states were correctly estimated are plotted. The data size of each condition is set to 100 *× (the numbe*r of states). For clarity, error bar is omitted because it should follow binominal distribution.

**Fig. S2.**
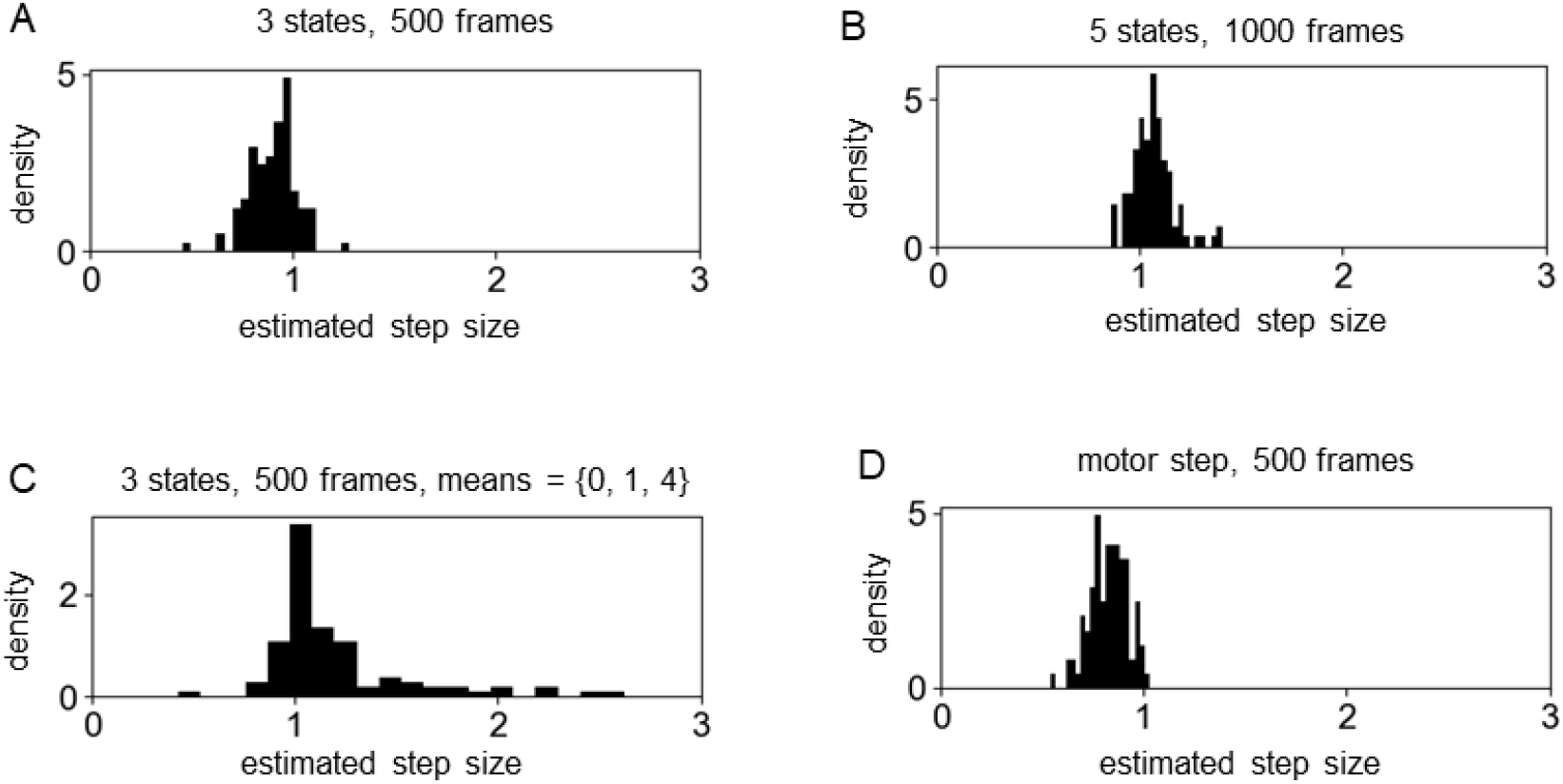
Histogram of estimated representative step size using the step finding results. Simulations were run 100 times for each condition. In all cases, the standard deviation of noise is set to 0.5. (A) three-state, 500-frame datasets were generated using mean values {1, 2, 3}. (B) five-state, 1000-frame datasets were generated using mean values {1, 2, 3, 4, 5}. (C) three-state, 500-frame datasets were generated using mean values {0, 1, 4}. (D) motor stepping trajectories were generated using forward step probability 0.03 and backward step probability 0.01.

**Fig. S3.**
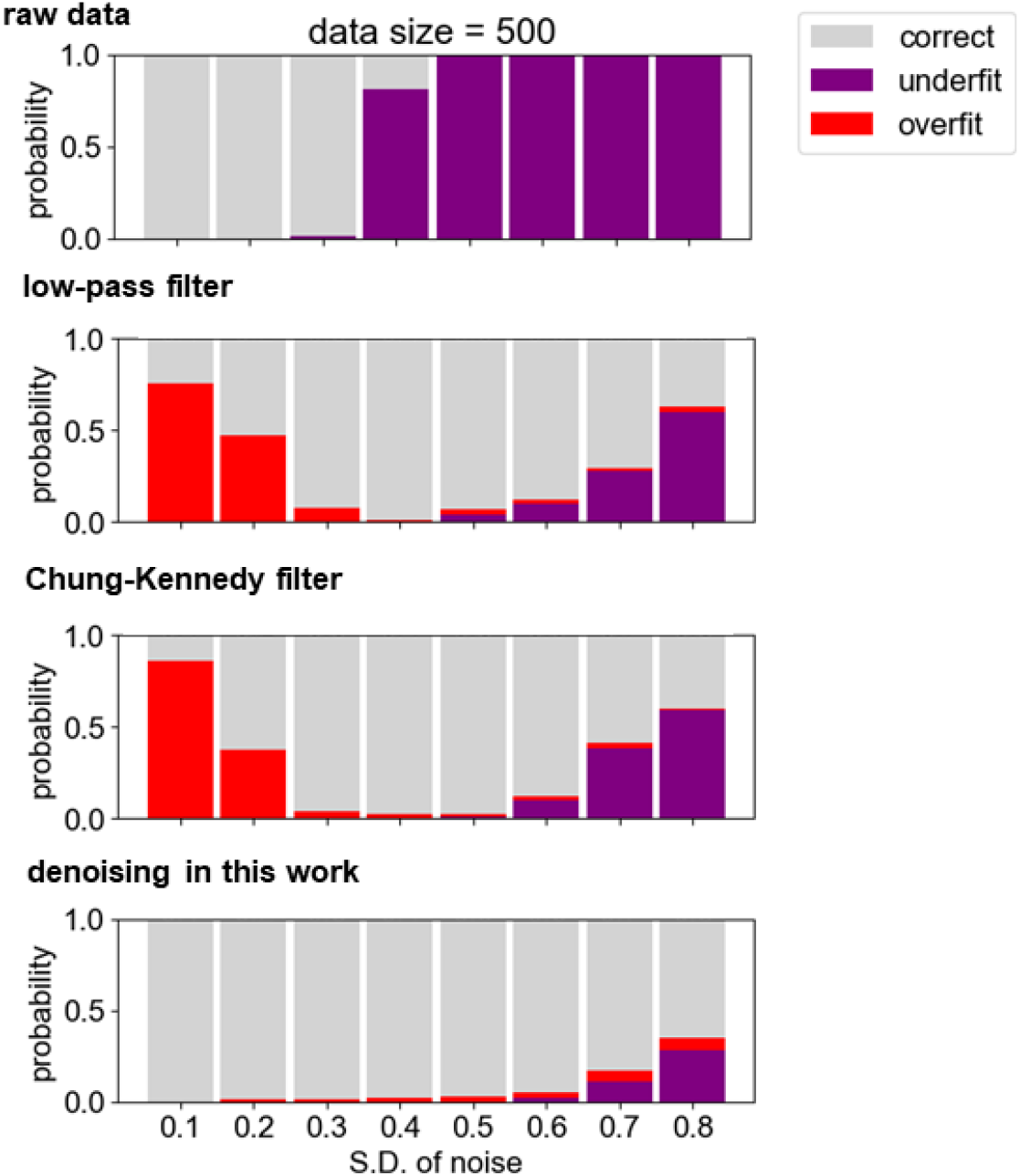
Probabilities of over-estimation (red) and under-estimation (purple) of the number of hidden states by GMM clustering, using raw data, low-pass filter, Chung-Kennedy filter and the denoising method in this work. For each condition, 3-state 500-frame sample datasets were generated similarly as Fig. 2 for 1000 times. User dependent parameters in low-pass filtering and Chung-Kennedy filtering are set to the same values as Fig. 3.

**Fig. S4.**
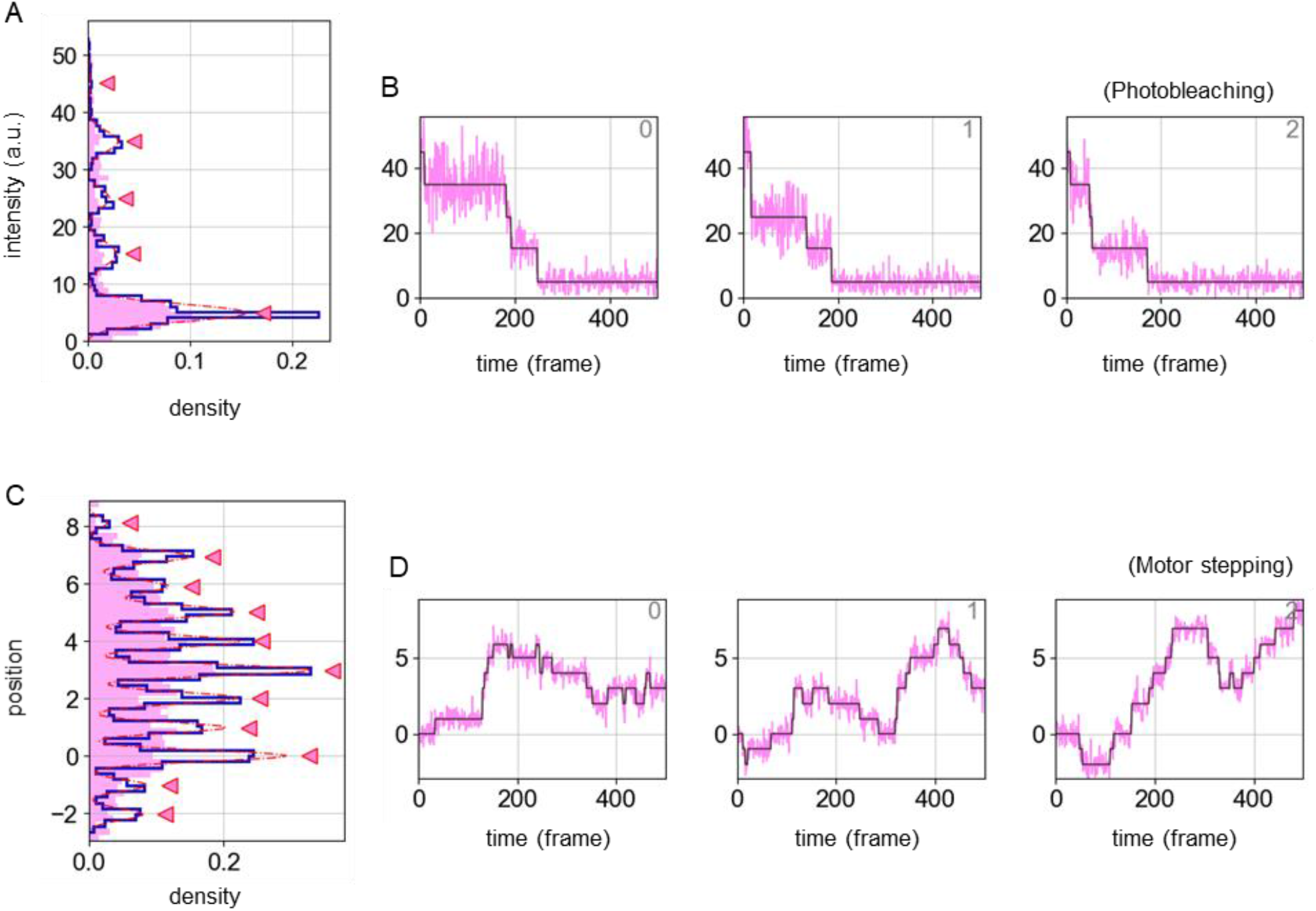
Application of sfHMM to HMM-type trajectories with sparse transition probability matrix. (A, B) Three trajectories are generated using mean values {5, 15, 25, 35, 45} with Poisson distributed signals, and fitted by sfHMM using Gaussian distribution model and default *p*_*sf*_. We did not fit with Poisson distribution model because photon counts may be unknown. (C, D) Three motor stepping trajectories are generated, and fitted by sfHMM using Gaussian distribution model and default *p*_*sf*_.

**Fig. S5.**
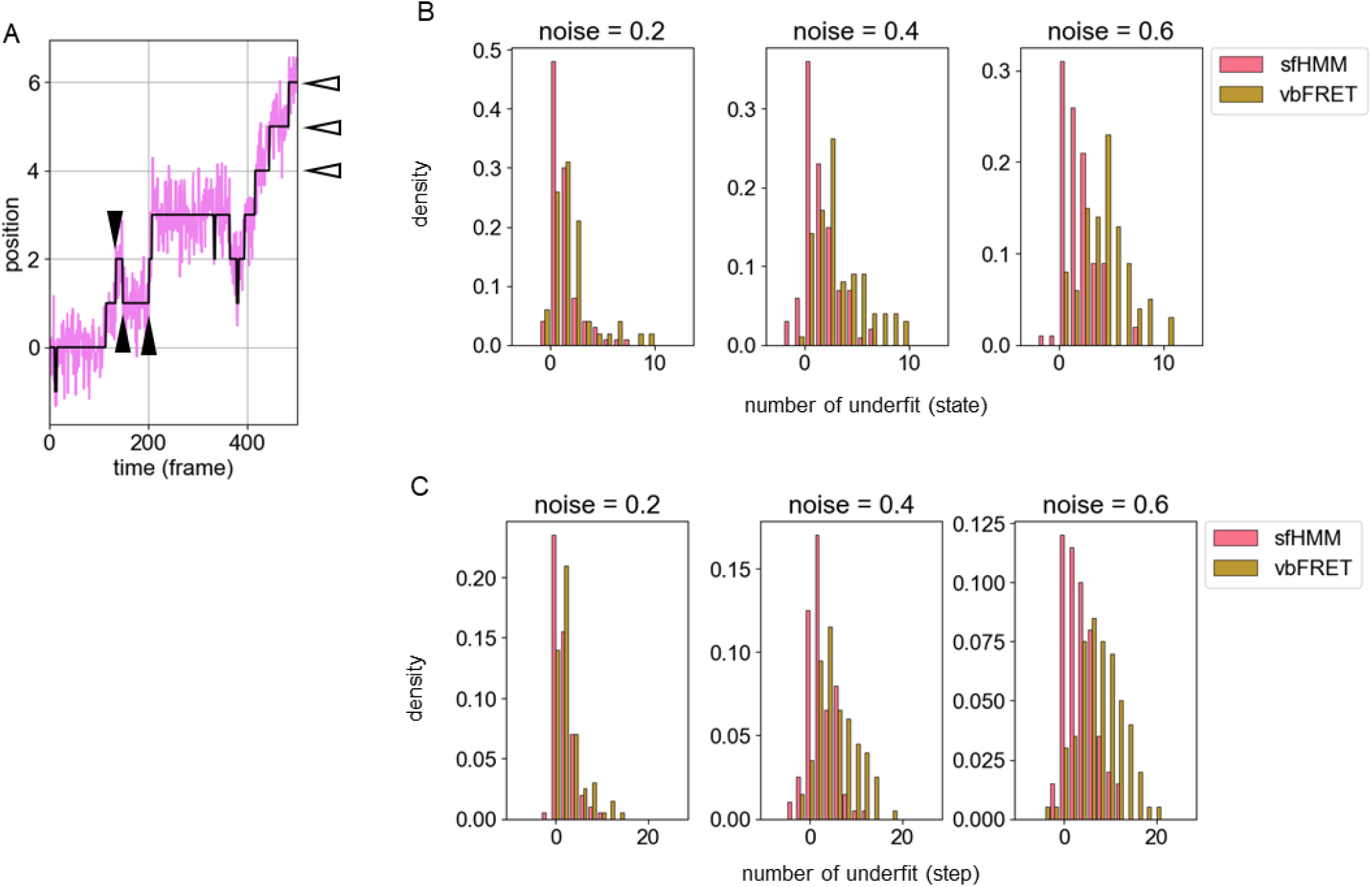
The results of applying sfHMM and _VB_FRET to motor stepping. 100 trajectories are generated using forward step probability 0.02 and backward step probability 0.01. (A) Analysis method. The white arrows indicate estimated “states” and the black arrows indicate estimated “steps”. (B) Histogram of estimated number of states with bin width 1. When a trajectory has *n*_*r*_ states and is estimated to have *n*_*e*_ states, then the number of underfit is defined as *n*_*r*_ *− n*_*e*_. Therefore, a negative value indicates overfit. (C) Histogram of estimated number of steps with bin width 3. The number of underfit is defined in a same way as in (B).

**Fig. S6.**
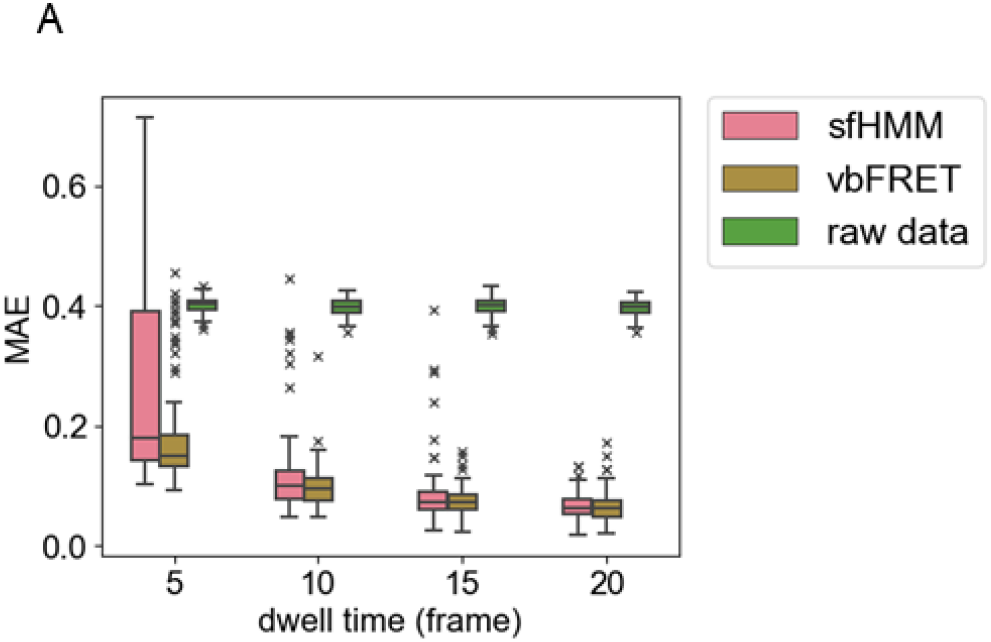
Comparison between sfHMM and _VB_FRET, using three-state HMM with transition probability matrix satisfying *W*_*ii*_ = 1 − *q* and *W*_*ij*_ = q/2. For each average dwell times, *q* is calculated as *q* = 1 − exp(−1/*τ*), like Fig. 4. 500-frame HMM data were sampled 100 times. Average transition probability *p*_*sf*_ is set to 0.08 for *τ* = 5, 10 and to 0.04 otherwise.

**Fig. S7.**
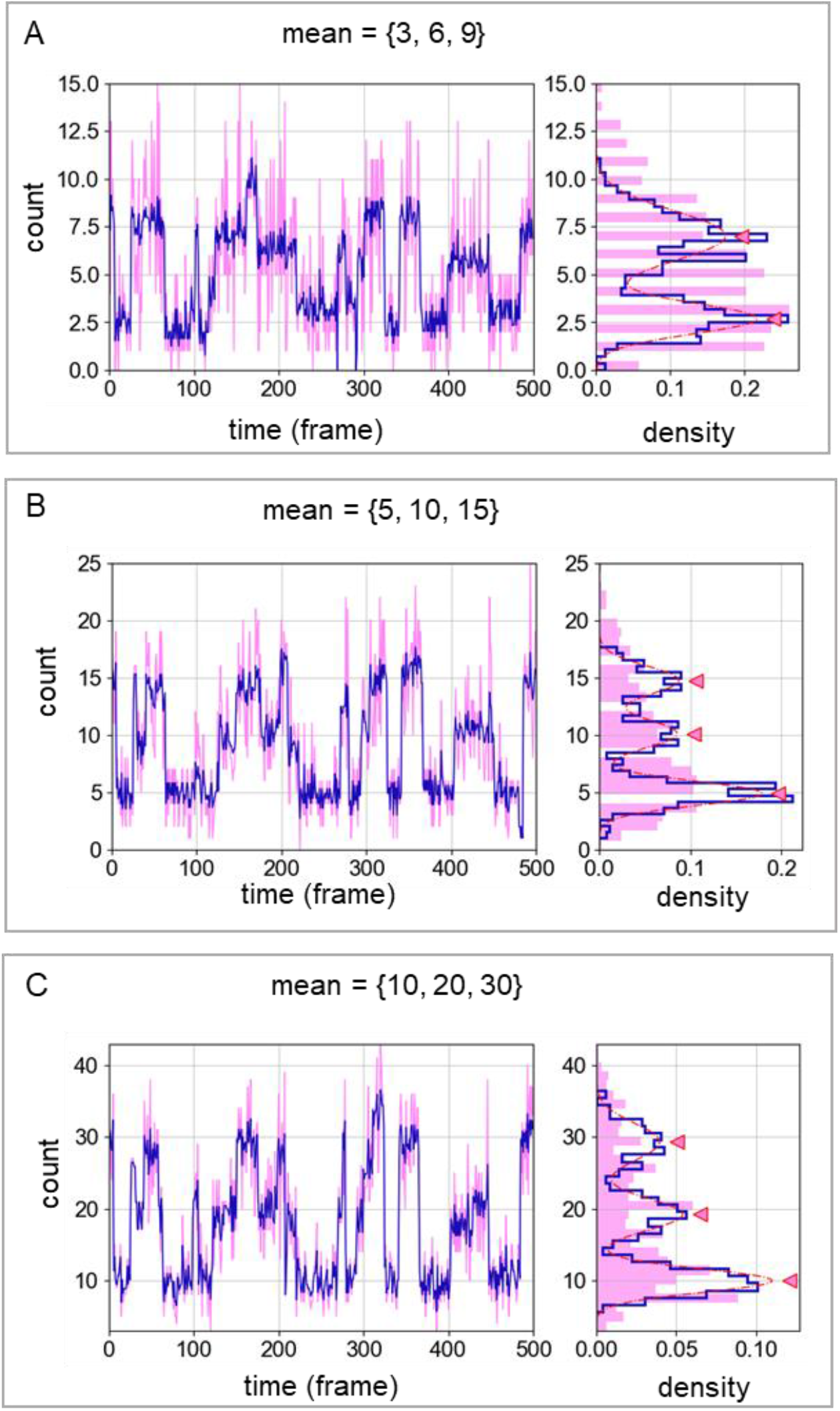
The GMM clustering results in sfHMM with different scales of Poisson distribution. Because the random seed was fixed, all the state transitions occurred at the same time for each condition.

## Appendix A. Maximum-likelihood-based step finding model

Given the observables ***d*** = {*d*_*1*_, *…, d*_*N*_} and the hidden states ***θ*** = {*θ*_*1*_, *…, θ*_*N*_}, according to the model proposed by Kalafut *et al*., likelihood is written as:

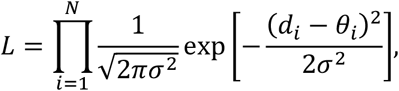

where *σ* is the standard deviation of Gaussian white noise. By solving *∂L/∂σ*^*2*^ = 0, the maximum likelihood estimator for *σ*^*2*^ is obtained as:

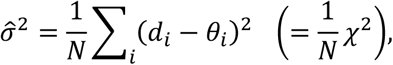

and the log-likelihood can be written as:

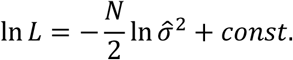

The last constant term is only dependent on the number of data ***d***. In the original algorithm, each step is considered as one free parameter, and the correct number of steps *k* are determined by BIC minimization:

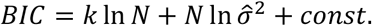

On the contrary, we introduced average transition probability *p*_*sf*_, assuming that whether the hidden state change occurs is determined probabilistically. This time, when hidden state changed *k* times and did not change *N* − *k* − 1 times, likelihood is written as:

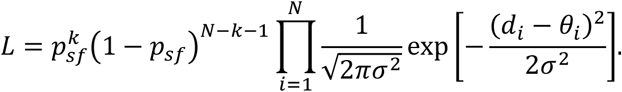

The maximum likelihood estimator 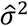 is identical to that shown above, and the log-likelihood is:

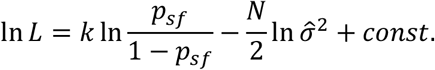

It turned out that BIC minimization in the original model is equivalent to likelihood maximization in our model, when 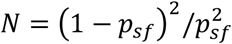 is satisfied. Therefore, our model is a natural extension of the original model.

A previous work proposed to multiply additional penalty factor *PF* to the BIC formula to avoid overfitting (Chistol *et al*., 2012). Our model is also interconvertible with that model by following equation:

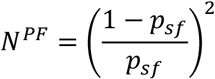

Parameter *PF* was an empirical one and needed experimental tuning. By taking the equation into consideration, we can translate *PF* into transition probability, which is easier to interpret.

## Appendix B. Derivation of log-likelihood for Poisson distributed signals

Poisson distribution is written as:

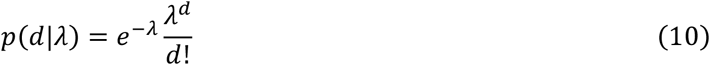

Assuming the hidden state changes at ***j*** = {*j*_*0*_ = 0, *j*_1_, ⋯, *j*_*k*_, *j*_*k*+1_ = *N* + 1}, then the maximum likelihood estimator for the parameter between *j*_*i*_ and *j*_*i*+1_ is:

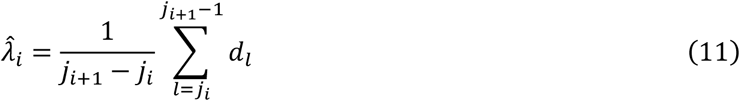

The log likelihood function can be calculated as:

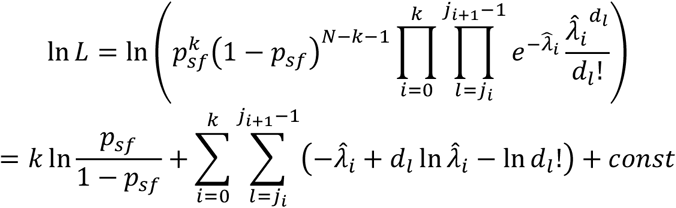

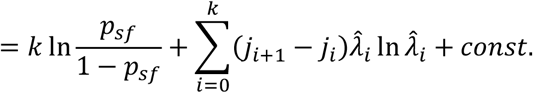

Unlike the log-likelihood function for Gaussian noise, the second term is independent of estimators in other steps. Therefore, all the steps can be determined in a faster way by applying greed algorithm (Ensign *et al*., 2010).

